# Auditory discrimination and identification of time of day for natural soundscapes

**DOI:** 10.1101/2025.08.28.672935

**Authors:** Frédéric Apoux, Nicole Miller-Viacava, Brian C.J. Moore, Bernie Krause, Jérôme Sueur, Christian Lorenzi

## Abstract

Following research on the auditory discrimination of natural soundscapes by human listeners [Apoux, Miller-Viacava, Ferrière, Dai, Krause, Sueur and Lorenzi (2023). J. Acoust. Soc. Am. 153, 2706], the present study assessed the ability of normal-hearing listeners to identify and/or discriminate the time of day (Dawn, Midday, Dusk, and Night) for soundscapes recorded in temperate and tropical forests. Identification was assessed using a single-interval four-alternative forced-choice task. Discrimination was assessed using a three-interval oddity task, where the listener was required to pick the odd one out. While identification of time of day was generally poor, discrimination was significantly above chance, indicating that listeners could hear some differences associated with the time of day, but could not effectively use those differences to assign the correct verbal label. The contribution of level, fine spectral, temporal envelope and temporal fine structure cues to discrimination was assessed. None of these cues appeared to be critical for time of day discrimination. Instead, it is suggested that listeners based their decisions on gross spectral cues, including aspects such as those related to ambient-noise spectral distributions.

## I. INTRODUCTION

Over the past few decades, the development of relatively new scientific fields such as “soundscape ecology” and “ecoacoustics” (for reviews, see Pijanowski *et al*., 2011; Sueur and Farina, 2015; Krause, 2016; Farina and Gage, 2017; Pijanowski, 2024) has sparked growing interest in the concept of the “soundscape”: the collection of sounds emanating from a given landscape. These sounds include those from biological (e.g., bird vocalizations), geophysical (e.g., wind and rain) and anthropogenic sources – those caused by humans (e.g., machinery). However, such a rudimentary definition of soundscape is not satisfactory for characterizing soundscapes operationally. In particular, several levels of soundscape integration may be considered, from the physical spatial and temporal patterns, i.e., the “distal” soundscape, to the subjective representation built by an observer, i.e., the “perceptual” soundscape (see Grinfeder *et al*., 2022). In the present work, the concept of perceptual soundscape is adopted as the operational definition. This refers to the perceptual auditory scene constructed from mixture of sounds reaching the ears from the distinct sources and how these sounds are shaped by the properties of the surroundings relative to a particular observer. This implies a subjective interpretation of the propagated sound signals. The present work focuses on “natural” soundscapes, i.e., soundscapes that contain as few anthropogenic sounds as possible.

Natural soundscapes are highly variable. They reflect a combination of ecosystem processes varying in space and time. Variations in space may be associated with changes in landscape patterns, biodiversity, and various physical factors such as temperature, solar radiation, elevation, and relative humidity (Sueur and Farina, 2015). When combined and further shaped by the sound propagation characteristics of the habitat, the acoustical patterns produced by these processes create soundscapes that are unique to that habitat. A few studies have shown that these unique patterns may be used to identify or discriminate soundscapes based on recordings made in specific locations. In a modelling study, Thoret *et al*. (2020) trained a support-vector machine to classify audio recordings from four habitats collected in the temperate terrestrial biome of the USA Sequoia National Park (Krause *et al*., 2011). As input to their model the authors used amplitude-modulation (AM) information extracted from the output of a model of the human auditory system composed of a cochlear filterbank followed by a modulation filterbank (Varnet *et al*., 2017). Classification scores were double the chance level, suggesting that spectro-temporal modulation cues can support soundscape identification. Following on from this, Apoux *et al*. (2023) asked human participants to discriminate between the four habitats considered by Thoret *et al*. (2020). Consistent with the modelling predictions, human listeners showed high consistency and high sensitivity when discriminating habitats based on short samples of natural soundscapes. However, it should be noted that Apoux *et al*. (2023) did not assess the identification of soundscapes; rather they assessed the ability to detect differences between soundscapes, based on any available cue.

As mentioned above, the acoustical patterns associated with a given landscape may vary over time. While some variations may be due to almost random factors (e.g., wind, rain), others reflect periodic events. For instance, the double-peak circadian pattern of animal vocal activity, the well-known dawn and dusk choruses, is associated with changes in spectro-temporal modulation information throughout the day (Gil and Llusia, 2020; Thoret *et al*., 2020). Such periodic occurrences may be used by human listeners to monitor the time of day (Lorenzi *et al*., 2023). Gage and Axel (2014) examined the temporal patterns of audio recording from an uninhabited island located on Twin Lakes in Grant Township, in Michigan’s northern Lower Peninsula (USA). They computed the power in each 1-kHz frequency interval between 1 and 11 kHz using an approach based on the average Power Spectral Density (PSD; Welch, 1967). Soundscape power in the 0-1 kHz interval was not reported as the authors pointed out that it is largely dominated by wind noise. Gage and Axel had recordings from six different sites on the island. They observed very little variation across sites, indicating that circadian patterns are very consistent across nearby locations. Therefore, they averaged the data over all sites and over a four-year period. The circadian patterns exhibited clear variations throughout the day, including multiple sharp peaks in different frequency bands. Despite the island itself being uninhabited, the 1-2 kHz band showed a broad peak during the day culminating in late afternoon around 16:00 and corresponding to anthropogenic activities on the lakes and on the lake shore. It also showed a smaller peak between approximately 02:00 and 04:00, associated with vocalizations of amphibians and common loons. The 2-3 kHz band showed a peak at 06:00 associated with the dawn chorus and a smaller peak around 20:00 associated with the dusk chorus. The same pattern was observed for the 4-5 and 5-6 kHz bands. The 3-4 kHz band showed a sharp increase in power shortly after sunrise, reaching a maximum between 06:00 and 08:00 and declining steadily over the day until about 20:00. A similar pattern was observed for the 6-7 kHz band. This power analysis illustrates that soundscapes are specific to a given time of day, and that the patterns of power variation across bands repeat from day to day, producing diel patterns that are consistent across years and nearby locations. It also illustrates how diel patterns are tightly coupled to the dawn and dusk choruses, producing a clear double-peaked pattern for the frequency bands 2-3, 4-5, and 5-6 kHz.

Other studies, however, show diel patterns that are site dependent. For instance, Loo *et al*. (2025) analyzed the daily and monthly temporal variations in soundscapes of a Malaysian rainforest using several ecoacoustic indices (for a brief description of each index, see Table I in Loo *et al*., 2025; for a review see Alcocer *et al*., 2022). The results varied with the index used but also with the recording site. Despite the variability across sites, the median hourly temporal variation of ecoacoustic indices generally showed either a simple bell curve with a peak (or dip) around noon or a flat pattern. The single peak centered around 12:00 contrasts with the double-peak circadian pattern found by Gage and Axel (2014). It should be noted that the ecoacoustic indices are typically derived from a selection of frequency bands and do not represent the power in each band. In any case, none of the indices used by Loo *et al*. (2025) clearly showed the effect of the dusk and dawn choruses.

Despite these discrepancies, it seems plausible that human listeners should be able to categorize soundscapes based on their time of recording. For the Twin Lakes database, this may be achieved by monitoring soundscape power information for multiple frequency bands in the 1-11 kHz region. Apoux *et al*. (2023) evaluated the ability of human listeners to discriminate soundscapes based on differences in time of day. In contrast to the hourly analysis of Gage and Axel (2014), Apoux *et al*. (2023) used four broad time periods. The rationale for using a limited number of broadly defined periods was that variations in soundscape mostly reflect the double-peaked circadian pattern of animal vocal activity and therefore only three to four periods may be clearly identifiable. A three-interval oddity task was used, the participants being required to pick the odd one out. The results showed that participants could discriminate soundscapes based on differences in time of day well above the chance score of 33%, scores ranging from 55 to 68% correct. However, Apoux *et al*. (2023) did not assess the absolute identification of time of day.

Several studies have assessed the sensitivity of human listeners to seasonality. Apoux *et al*. (2023), using the same approach as for assessing discrimination of time of day, reported above-chance scores ranging from 52 to 66% correct for season discrimination (spring, summer, autumn, winter). Again, they did not assess absolute identification. McMullin *et al*. (2024) asked human listeners to judge four different global properties of 200 scenes. The scenes were 4-s long, and listeners were allowed to listen to each scene as many times as they needed to make each of the four judgments. One property was season (“What season does this scene sound like?”). The authors used a seven-point Likert scale ranging from 1 to 7 (highest extreme) that included the four seasons but also three “between seasons” categories that are not naturally used: Between Winter and Spring, Between Spring and Summer and Between Summer and

Fall. In contrast to Apoux *et al*., McMullin *et al*. (2024) asked the participants to directly identify the season. While accuracy was low at only 25.3% correct, it was well above the chance level of 14.3%. Moreover, some design choices may have artificially increased the difficulty of the identification task and therefore lowered the scores. First, the decision to include intermediate seasons may have been slightly confusing for the participants as these intermediate seasons are not very well defined in the popular imagination and rarely used in practice. Second, the stimuli used by McMullin and colleagues consisted of 200 auditory scenes from various locations across the United States. According to the authors, they were chosen to “represent typical environments humans are exposed to, such as parks, classrooms, hiking trails, city streets, forests, and cafes.” This large diversity had several potential consequences for the ability to identify seasons. Because the database included multiple recordings from similar but distinct settings (e.g., different cafés) and they did not record each season for each location, participants might have had to identify Winter in a café in Los Angeles and Spring in a classroom in Miami. This lack of consistency most likely reduced the ability to establish the cues associated with each season. Another consequence is that several recordings were made indoors. It is likely that these settings did not convey as much information about the season as outdoor settings, further complicating the extraction of cues associated with seasons from the samples. Despite these limitations, participants in McMullin *et al*. (2024) were able, to some extent, to identify seasons. Therefore, the above studies demonstrate both that human listeners are sensitive to the differences introduced by periodic variations in soundscapes, and that they are able, to some extent, to label scenes based on the time of recording.

The present study extended previous work by investigating the sensitivity of human listeners to diel acoustical variations in natural soundscapes. A first goal was to replicate the work of Apoux *et al*. (2023) using a comparable (i.e., a temperate forest) but distinct biome, so as to assess the generalizability of the initial results. A second goal was to evaluate the sensitivity of human listeners to time of day not only in terms of discrimination, but also in terms of absolute identification.

The analyses of variations in power in different frequency bands conducted by Gage and Axel (2014) showed that diel variations in a single area, the Twin Lakes, were largely driven by animal vocalizations. Based on this, Apoux *et al*. (2023) hypothesized that discrimination performance may improve as biotic activity and diversity increase. A third goal of the present study was to assess this assumption. While several tools have been developed to estimate the contribution of biophony to the acoustical properties of a given habitat (e.g., Farina and Gage, 2017; Grinfeder *et al*., 2022; Pijanowski, 2024), a general approach was adopted here. Several biomes varying in their expected biodiversity were selected, informed by the “latitudinal gradient of diversity”, namely that species numbers are maximal at low latitudes and decrease from the Equator towards either Pole. This latitudinal gradient has been shown for many groups of vertebrates and invertebrates, for terrestrial and aquatic organisms and for several continents (Willig *et al*., 2003). Importantly, it is especially striking for birds (Gaston, 2000; Hillebrand, 2004; for a review, see Rolland and Freeman, 2022). Consequently, if biophony plays a significant role in the discrimination of times of day, performance should be better for biomes at low latitudes (with high species richness), than for biomes at high latitudes (with low species richness).

A fourth goal of the present study was to investigate the acoustic cues involved in time of day discrimination. Apoux *et al*. (2023) conducted a similar investigation for habitat discrimination and found that human listeners seem to base their decisions on gross spectral cues (timbre) related to biological activity and/or habitat acoustics, with greater weight for frequency components in the mid-frequency range (1-3 kHz). Although they also assessed time of day discrimination, they did not address the issue of the cues used to perform this task. To examine this, the approach used by Apoux *et al*. (2023) was implemented: stimuli were restricted in the audio-frequency domain or were noise vocoded to assess the contribution of global spectral cues and temporal-envelope cues, respectively.

## II. GENERAL METHODS

### A. Participants

Twenty-five normal-hearing listeners (15 females) participated in the experiments. Normal hearing was defined as passing a pure-tone air-conduction screening at 20 dB HL for octave frequencies from 250 to 8000 Hz, measured on the first day of testing. All listeners were recruited through the platform RISC (*“Relais d’Information sur les Sciences de la Cognition, UMS CNRS 332”*) at Ecole normale supérieure (Paris, France). The listeners were fully informed about the purpose of the study and provided written consent before their participation. All listeners were paid 10 Euros per hour. This study was approved by the national ethical committee CPP Ile de France III (Am8618-1-S.C.3460; N° EUDRACT: 2016-A01769-42).

### B. Stimuli

The stimuli were generated from three sound databases. Figure 1 shows a picture of each recording location.

**Figure 1:**
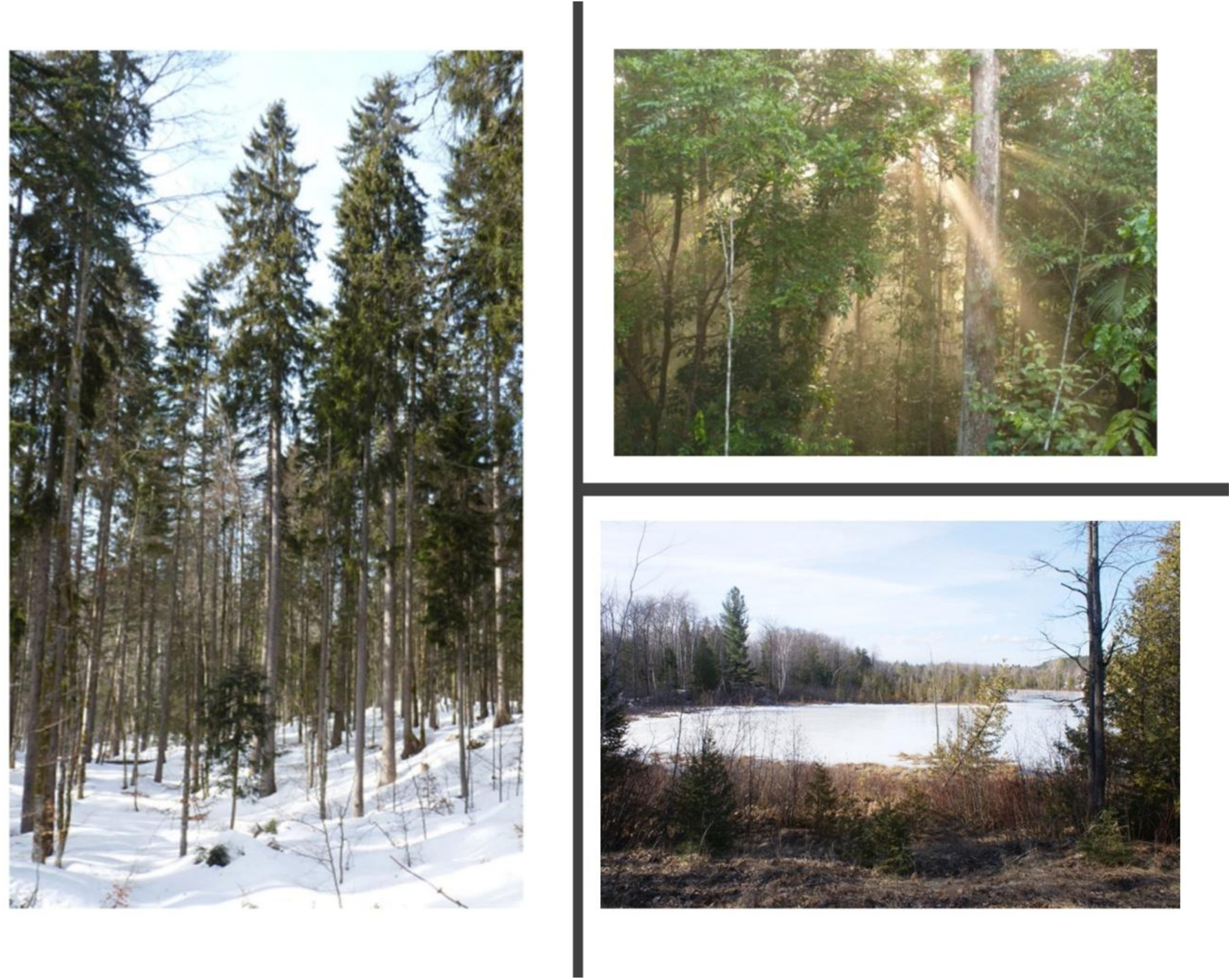
Pictures of the three recording locations employed in the experiments. Risoux, Haut Jura, France (left); CNRS Research Site, Nouragues, French Guyana (top right); Twin Lakes, Michigan, USA (bottom right).

The first database, obtained at Twin Lakes (TL), was used in Experiments I and II. It was recorded by Gage and Axel (2014) at the water-vegetation edge of an uninhabited island located on Twin Lakes in Grant Township, in Michigan’s northern Lower Peninsula, USA (average elevation of 210 m). This is a forested island with a mixture of deciduous and coniferous vegetation, including birch, trembling aspen, balsam fir, white cedar, tamarack and white pine. One-minute recordings were made every 30 minutes at six sites on the island over a four-year period (2009-2012) from April to October for a total of 48 samples per day/site.

The second database, Haut Jura (HJ), was used in Experiment II only. It was collected by the Museum national d’Histoire naturelle (MNHN) team (Grinfeder *et al*., 2022) in the Risoux forest. This 4400-ha cold coniferous forest is located in the Jura mountains, in the east of France. The site is mainly covered by spruce *Picea abies* forest. The habitat structure undergoes small changes over the year, with snow cover during the winter and slight changes of the foliage density due to the presence of some deciduous trees and shrubs. The average elevation is 1230 m. One-minute recordings were made every 15 minutes at multiple sites across the forest all year round over a one-year period from August 2018 to July 2019, for a total of 96 samples per day/site.

The third database, Nouragues (No; de Baudouin *et al*., 2023), was used in Experiment II only. The site is a neotropical lowland dense evergeen rainforest (French Guiana, France), near the Saut Pararé rapids of the Arataye River (CNRS Nouragues Research Station). There are two main seasons, a dry season from September to December and a rainy season from January to June. A secondary short dry season occurs in March within the rainy season. The forest is characterized by a fairly open understorey but with a dense canopy between 40 and 45 m. The ground is covered with thick litter. The average elevation is about 100 m. One-minute recordings were made every 15 minutes at a single site (4°2′N; 52°40′W), starting February 2019 (still ongoing), with a total of 96 samples per day.

One set of stimuli was extracted from each database using the following procedure. First, a single year and recording site were selected for each location. To provide a representative sampling of the changes occurring throughout the year (due to biological activity and climatic variations), three months were chosen based on average precipitation. This factor was preferred over season as it was not possible to define season consistently across databases. The three months were selected independently for each location and corresponded to the months with the lowest, median, and highest probability of precipitation. These months were April, July and October for TL, March, December and June for HJ, and September, December and May for No, all in increasing order of probability of precipitation.

Five evenly spaced days were selected from each month for a total of 15 days per location. The days were then divided into four two-hours periods corresponding to four times of day: Dawn, Midday, Dusk and Midnight. First, sunrise and sunset were computed for each day in order to reflect real sunlight^1^. Then, Midday was defined as the two hours centered around the middle point between sunrise and sunset. Midnight was defined as the two hours centered around the middle point between sunset and sunrise with the constraint that it should always be fully included in the selected day. All times were rounded up or down to the nearest hour. Finally, nine 1-s samples evenly spaced within each two-hour window were extracted, resulting in a set of 540 samples for each biome (3 months × 5 days × 4 times of day × 9 samples). The onsets and offsets of all samples were tapered using 50-ms raised-cosine ramps. Natural disparities in level across samples were preserved for each dataset. All stimuli were presented diotically at an average level of 55 dB SPL(A), using Sennheiser HD650 circumaural headphones. This level was calculated for each biome independently so that the average level of each dataset was 55 dB SPL(A).

Figure 2 shows average two-dimensional amplitude-modulation (2D-AMi) spectra for each time of day, computed using a model of human auditory processing (Thoret *et al*., 2020; Varnet *et al*., 2017) for each database. The samples were first passed through a bank of bandpass filters tuned in the audio-frequency domain that simulated peripheral (cochlear) filtering (70-11025 Hz) (Glasberg and Moore, 1990). The temporal envelope at the output of each simulated cochlear filter was then extracted and passed through a second bank of bandpass filters simulating modulation filtering by humans (0.5-200 Hz, Q = 1) (Moore *et al*., 2009). The 2D-AMi spectra in Fig. 2 simulate the differences in AM information across biomes and across times of day available to the central auditory system of humans. As can be seen, each biome exhibits a distinctive pattern. For each biome, there are also subtle changes with the time of day. For HJ, Midnight was characterized by higher energy for low spectral channels and high modulation rates. For No, the 2D-AMi spectra revealed a difference between Midnight and other times, with more energy for Midnight for the low-to-mid spectral channels and low-to-mid modulation rates. For TL, there was a difference between Dawn and other times, with more energy for the high spectral channels and low-to-mid modulation rates.

**Figure 2:**
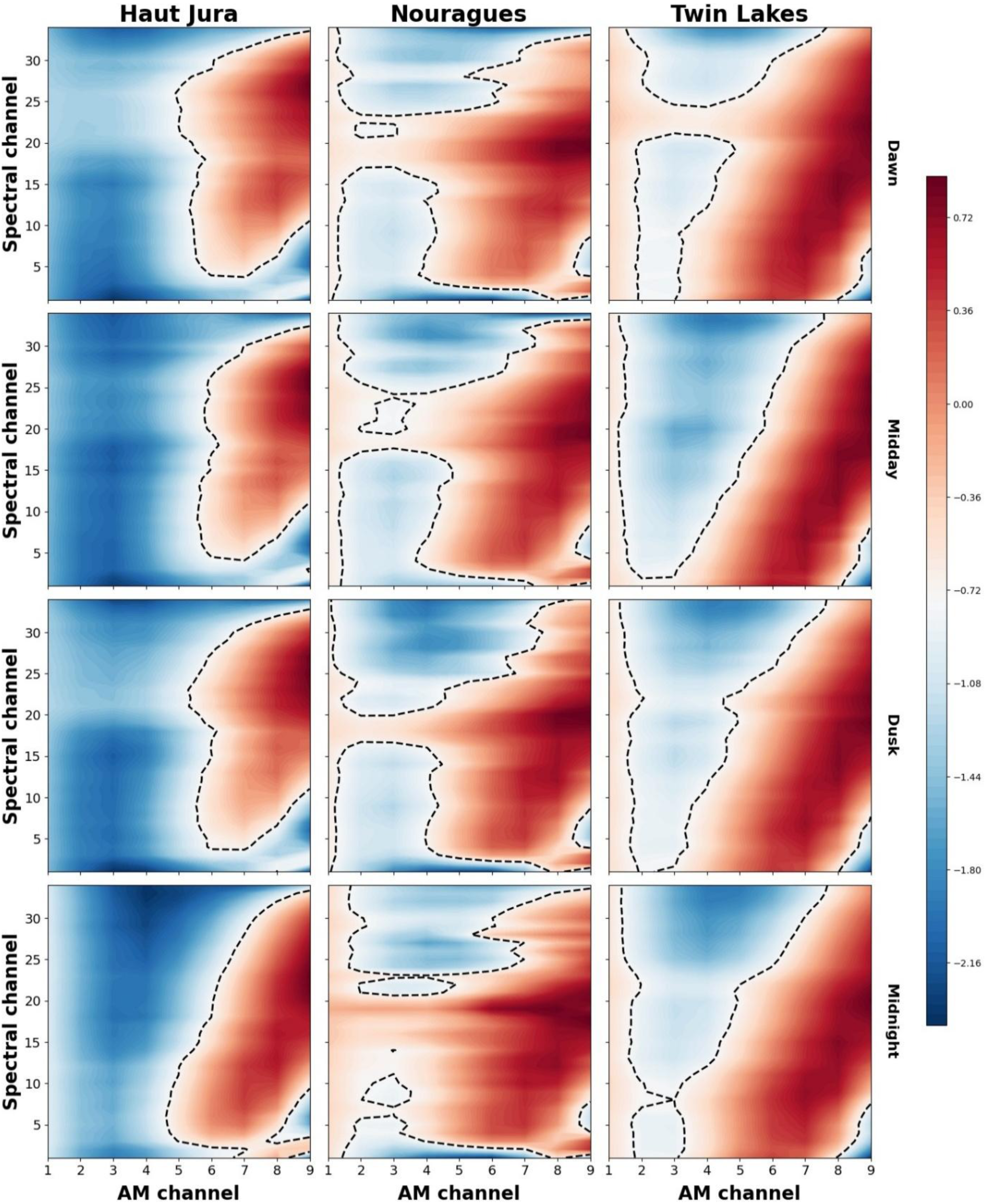
Two-dimensional amplitude-modulation (2D-AMi) spectra computed using a model of the human auditory system for the three natural soundscapes databases recorded at dawn, midday, dusk and midnight. The 2D-AMi spectra show the distribution of AM energy (as computed by the modulation index, AMi) plotted in dB as a function of AM (abscissa) and audio frequency (ordinate) channel for each biome and time of day, where the color indicates the range of values from high (red) to low (blue). The dashed lines indicate the midrange energy value. From left to right, temperate forest (Haut Jura, France), tropical forest (Nouragues, French Guyana), temperate forest (Twin Lakes, USA). From top to bottom, dawn, midday, dusk and midnight.

### C. Procedure

Two psychophysical paradigms were implemented to assess the ability to identify and discriminate times of day. Because the multiple-interval oddity paradigm may have lower ecological validity (Schmuckler, 2001; Neuhoff, 2004; Keidser et al., 2020) than the identification paradigms typically used in environmental studies (e.g., Gygi *et al*., 2004; Shafiro, 2008), identification tasks were included in the present work. A single-interval, four-alternative procedure and the method of constant stimuli was used to assess identification. On each trial, the participant was presented with a 1-s sound sample and asked to identify the time of day from four possible choices (Dawn, Midday, Dusk and Midnight) by clicking on one of four boxes presented on a computer screen. All 540 samples were presented during a test session (a block), resulting in 540 trials. Samples were randomly selected without replacement within a test session, so that no stimulus was presented more than once. Visual feedback was given at the end of each trial (see specific experiment for details of feedback). The feedback was displayed 0.5 s after a response was made and the next trial started 1 s after feedback was given. After every twenty trials, listeners were given a 10-s break. Each identification session lasted about 30 minutes.

A 3-interval forced-choice oddity paradigm (Frijters, 1979; Frijters *et al*., 1980; Versfeld *et al*., 1996) and the method of constant stimuli were used to assess discrimination. On each trial, three successive stimuli were presented, one target and two standards, separated by a 1-s silent interval. The three stimuli were extracted from recordings made on the same day, but the day could differ across trials. The interval containing the target was selected randomly. The task was to indicate the interval that differed from the other two intervals, by clicking on one of three boxes presented on a computer screen. Each test session consisted of 180 trials for a total of 3×180 = 540 unique stimuli per session. Samples were selected without replacement so that no stimulus was presented more than once during a session. Note that the two standards within a trial were not identical, although they were drawn from the same time of day. Thus, the task could not be performed reliability based on small spectro-temporal details of the stimuli, since these differed for the two standards within a trial. Visual feedback as to the correctness of the answer was given at the end of each trial. Feedback was provided 0.5 s after one of the response boxes was pressed, and the next trial started 1 s after the feedback was given. After every twenty trials, listeners were given a 10-s break. Each session lasted about 20 minutes.

## III. EXPERIMENT I

### A. Method

Ten normal-hearing listeners (five females) took part in Experiment I. However, the data for one participant were excluded for the following reasons. First, it was determined using the interquartile range (IQR) method that 50% of the data points from this participant could be considered outliers, with half of them being extreme outliers (3×IQR). Only two other data points from two different participants were outliers. Second, removing the data for this participant did not change the results of the statistical analysis but did affect the assumption of normality, the overall data being normally distributed only after the data from this participant were removed. Listeners were aged between 21 and 37 years (mean = 28 years; standard deviation, SD = 5.9 years). Experiment I included four tasks.

#### Ia. Identification task

Experiment Ia explored the ability to identify time of day using four broad time windows (Dawn, Midday, Dusk and Midnight). Simple “right” or “wrong” feedback was provided; the correct answer was not disclosed.

#### Ib. Discrimination task: spectral filtering

Experiment Ib explored the contribution of different spectral regions to the ability to discriminate time of day. In one condition, the stimuli were intact, i.e., they were not filtered (UNF). In eight other conditions, they were lowpass (LP) or highpass (HP) filtered (zero phase, –72 dB/oct slope Butterworth filter) at each of four cutoff frequencies (0.5, 1, 2, and 4 kHz), giving nine conditions (including UNF). There were nine sessions, each corresponding to a given condition, which were tested in random order across participants. Experiment Ib lasted about 4 h (including breaks).

#### Ic. Discrimination task: noise vocoding and envelope filtering

Experiment Ic explored the contribution of temporal fine structure (TFS) and temporal-envelope cues to the ability to discriminate changes in time of day by noise vocoding the stimuli and manipulating the envelope fluctuations present in the signal. Noise vocoding replaces the original TFS in the signal with less informative TFS (Moore, 2014). Each waveform sample was initially filtered into eight, 4-ERB_N_-wide (Glasberg and Moore, 1990) non-overlapping frequency bands spanning the range 80–9500 Hz (zero phase Butterworth filters, 72 dB per octave roll-off). The nine cutoff frequencies were set to 80, 246, 502, 894, 1499, 2428, 3856, 6054, and 9433 Hz, resulting in center frequencies (CF) of 154, 360, 677, 1164, 1913, 3066, 4837, and 7562 Hz for bands 1 to 8, respectively. The Hilbert transform was applied to the output of each frequency band to decompose the bandpass-filtered signal into its temporal-envelope (modulus of the Hilbert analytic signal) and TFS (cosine of the argument of the Hilbert analytic signal). The resulting envelopes were LP or HP filtered (zero-phase Butterworth filter, 72 dB/oct roll-off) and then used to modulate Gaussian white noises. The modulated noises were frequency-limited by filtering with the same bandpass filter as used for the original analysis band and assigned their original long-term rms power to preserve the overall spectral shape. Finally, the resulting modulated noises were summed up. In one condition, the envelopes were lowpass filtered at half the bandwidth of the auditory filter at the CF of the analysis band (ERB_N_/2) and the original TFS was used as carrier. This “reconstructed” condition (REC) carried TFS information more like that for the original stimuli than for the noise-vocoded stimuli. In five other conditions, the envelopes were LP or HP filtered at each of two cutoff frequencies (5 and 20 Hz). A condition with the envelopes LP filtered at ERB_N_/2 was also used, giving a total of six vocoded conditions. Discrimination was tested in six sessions, one for each condition, with the order randomized across participants. Experiment Ic lasted about 3 h (including breaks).

#### Id. Discrimination task: equalized rms

Since it was decided to preserve natural level cues in the main experiments, an additional experiment was conducted to assess the contribution of these cues to the discrimination of changes in time of day. In Experiment Id, all stimuli were equated in terms of long-term rms power, to reduce loudness cues. Discrimination was tested in a single session, using intact stimuli. Experiment Id lasted about 20 minutes.

Participants completed the first three experiments in order Ia, Ib, and Ic. Experiment Id was completed after one of the other experiments, which of these being determined randomly for each participant.

### B. Results

#### Ia. Identification task

Identification scores for each participant are shown as “Session 1” in Fig. 8 (the remainder of Fig. 8 is discussed later). The scores ranged from 23 to 32% with a mean of 28% and a standard deviation of 3 percentage points. The mean score was slightly but significantly above the chance level of 25% (one-sample *t* test, *p*<0.01). However, it is apparent that none of the participants was able to reliably identify the time of the day for the TL biome.

#### Ib. Discrimination task: spectral filtering

Experiment Ib explored the contribution of different spectral regions to the ability to discriminate time of the day. Figure 3 shows individual discrimination scores for the LP (left panel) and the HP (right panel) conditions. In each panel, the dashed line indicates performance averaged across all nine participants and the dotted line indicates performance in condition UNF, which ranged from 50 to 67%. Performance was above chance for all conditions for each participant. There was variability in the pattern of results across participants, but the average data (dashed lines) show a monotonic effect of cutoff frequency, performance decreasing by about 4 percentage points with decreasing LP cutoff frequency and increasing by about 5 percentage points with increasing HP cutoff frequency. The two functions crossed slightly above 2 kHz.

**Figure 3:**
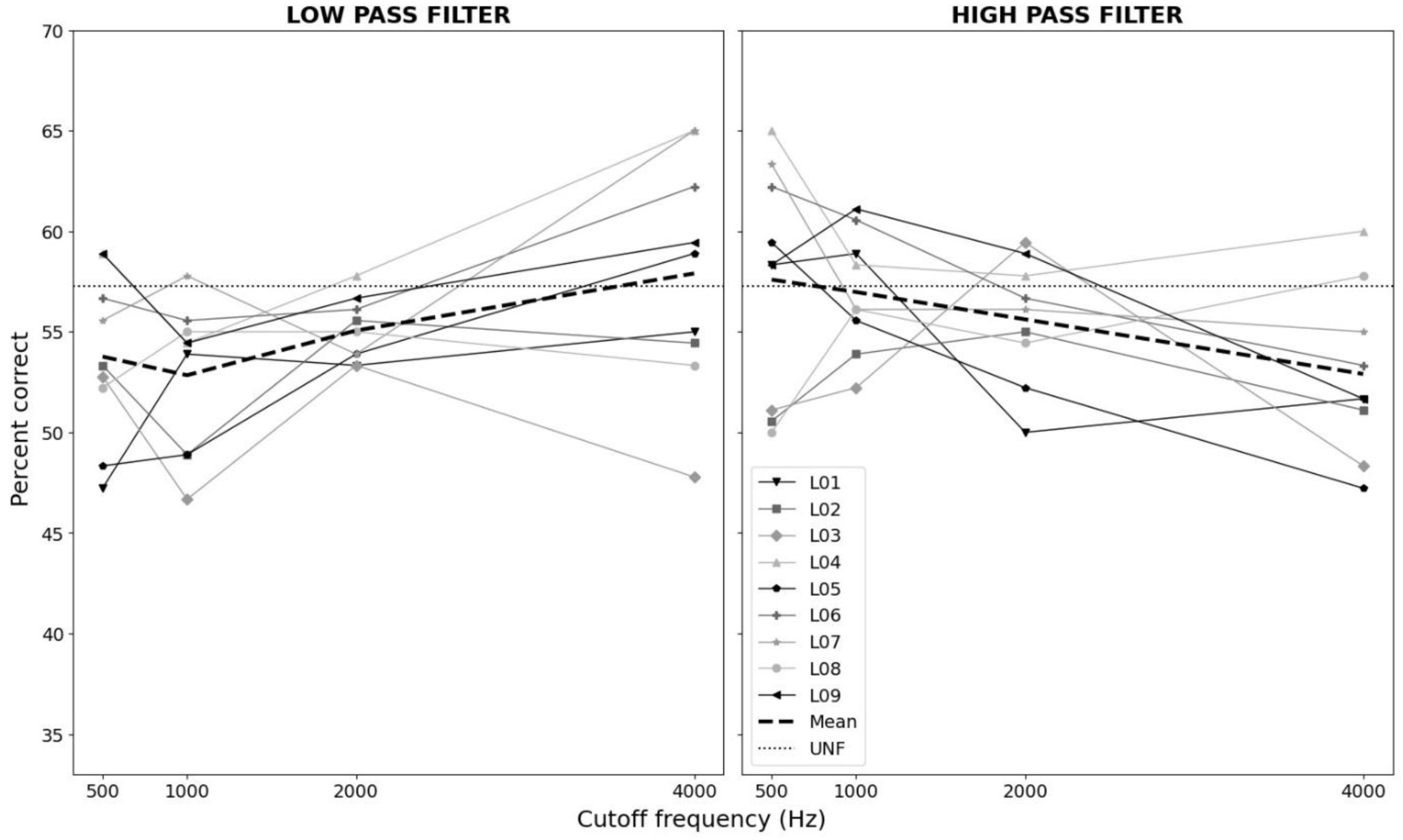
Individual data for Experiment Ib. Scores for discrimination of time of day are plotted as a function of cutoff frequency for stimuli that were either lowpass (left panel) or highpass filtered (right panel) in the audio-frequency domain. In each panel, the horizontal dotted line shows the score for the unfiltered condition (UNF); the dashed line shows scores averaged across all participants.

A repeated-measures analysis of variance (ANOVA) conducted on the discrimination scores with filter type (2 levels) and cutoff frequency (4 levels) as within-subjects factors revealed that the main effect of filter type [*F*(1,64)=0.82; *p*=0.369] and the main effect of cutoff frequency [*F*(3,64)=0.11; *p*=0.955] were not significant, but there was a significant interaction between filter type and cutoff frequency [*F*(3,64)=4.76; *p*<0.005], consistent with performance changing in opposite directions for the two filtering conditions with changing cutoff frequency.

To further investigate whether there was a significant relationship between cutoff frequency and performance, a linear regression analysis was performed for both filter types. After fitting the linear regression model, the slope was found to be significantly different from 0 for both LP (*p* < 0.05) and HP (*p* < 0.05), suggesting a significant linear relationship between cutoff frequency and performance. Overall, best discrimination was found for the unfiltered condition, but performance was above chance even when the stimuli were LP filtered at 500 Hz or HP filtered at 4000 Hz.

#### Ic. Discrimination task: envelope filtering

Experiment Ic explored the contribution of TFS and temporal-envelope cues to the ability to discriminate changes in time of the day. Figure 4 shows individual discrimination scores for the LP (left panel) and the HP (right panel) envelope-filtering conditions using noise-vocoded stimuli. In each panel, the dashed line indicates performance averaged across all nine participants and the dotted line indicates performance in condition REC, for which TFS information was represented more accurately than for the noise-vocoded stimuli. Performance in condition REC ranged from 47 to 63% and was well above chance for each participant. Performance was almost identical for condition UNF of experiment Ib (57%) and condition REC (58%). This suggests that condition REC did indeed preserve the important features of the original stimuli. Performance in the condition with envelope lowpass filtered at ERB_N_/2 (not shown) ranged from 44 to 61% (mean=56.7, SD=5.3).

**Figure 4:**
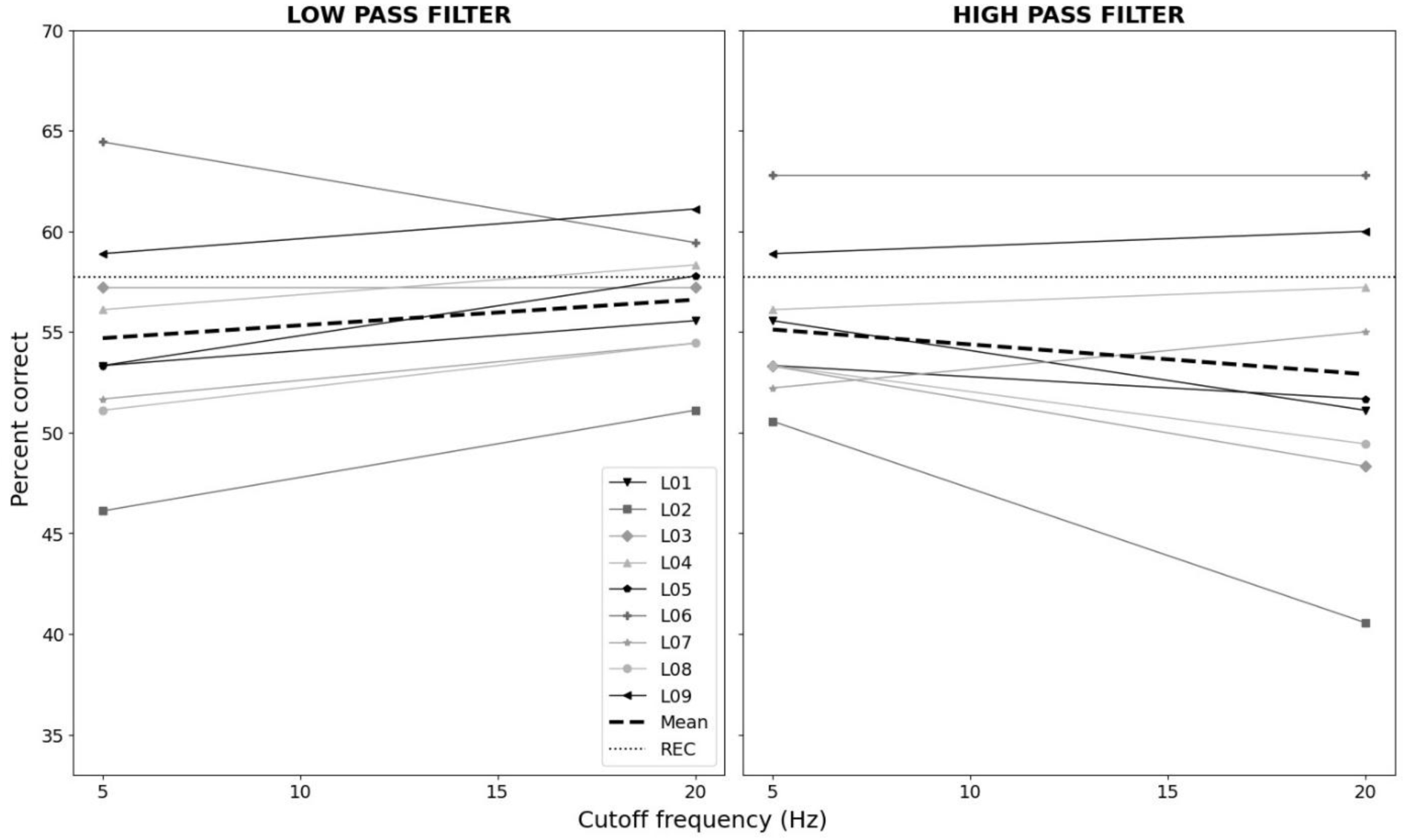
Individual data for Experiment Ic. Scores for discrimination of time of day are plotted as a function of envelope cutoff frequency when the envelopes of vocoded stimuli were either lowpass (left panel) or highpass filtered (right panel) in the time domain. In each panel, the horizontal dotted line shows the score for the reconstructed condition (REC); the dashed line shows the score averaged across all participants.

Performance was only a little worse for the noise-vocoded stimuli than for the REC stimuli, suggesting that near-intact TFS information and fine spectral detail were not critical for discrimination performance. Also, the effect of envelope filtering was small, performance dropping by only about 2 percentage points when the envelope LP cutoff frequency was decreased from 20 to 5 Hz and by about 4 percentage points when the envelope HP cutoff frequency was increased from 5 to 20 Hz.

A repeated-measures analysis of variance (ANOVA) of the discrimination scores with filter type (2 levels) and cutoff frequency (2 levels) as within-subjects factors showed no significant effects of filter type [*F*(1,32)=1.00; *p*=0.324] or cutoff frequency [*F*(1,32)=0.01; *p*=0.926]. The interaction between filter type and cutoff frequency was also not significant [*F*(1,32)=1.60; *p*=0.215].

#### Discrimination tasks: influence of time of day and season

The results of experiments Ib and Ic were examined further to assess the potential influence of the time of day and the season on performance. Scores were computed separately for each time of day (Fig. 5, left panel) and each season (Fig. 5, right panel) for the UNF and REC conditions only.

**Figure 5:**
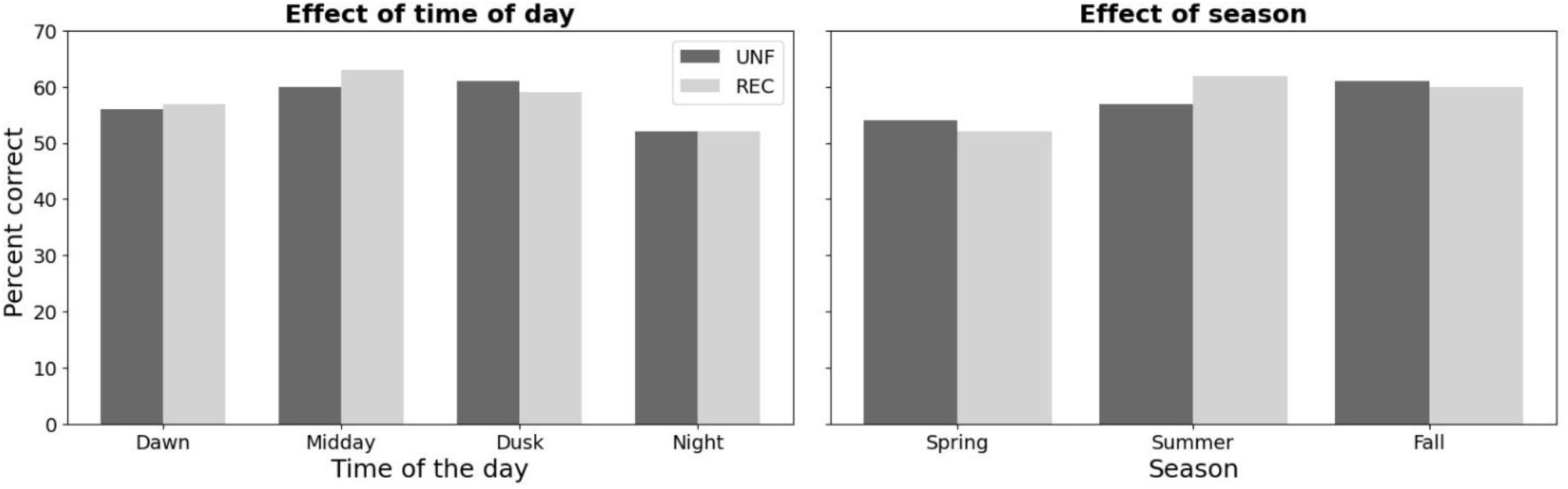
Mean data for Experiments Ib and Ic. Discrimination scores are plotted as a function of the time of day (left panel) or as a function of the season (right panel) for the unfiltered (UNF) and reconstructed (REC) conditions.

Performance was somewhat affected by the time of day, varying from 51% (Night) to 61% (Dusk) for UNF and from 50% (Night) to 61% (Midday) for REC. A repeated-measures ANOVA of the discrimination scores with processing type (2 levels) and time of day (4 levels) as within-subjects factors showed no significant effect of processing type [*F*(1,8)=0.09; *p*=0.773] and a significant effect of time of day [*F*(3,24)=6.93; *p*<0.05]. The interaction between processing type and time of day was not significant [*F*(3,24)=0.64; *p*=0.60]. These effects are difficult to interpret. Although biophony is typically lower during the night, it is generally associated with species that are less active during daytime and which are therefore somewhat identifiable. For instance, Gage and Axel (2014) reported that nighttime sounds in Twin Lakes may be attributed to insects, green frogs (*Rana clamitans*), northern spring peepers (*Pseudacris crucifer*), common loons (*Gavia immer*) and Canada geese (*Branta canadensis maxima*), some of these animals being active only during the night. It might have been assumed that participants would rely on sounds that are specific to nighttime to more accurately discriminate night from the other times of day. This was not the case for the present experiment. However, additional analyses revealed that performance was always lower when night was the standard. This was the case for both condition UNF and condition REC.

To further explore these unanticipated results, the energy in 10 frequency bands was computed, as in Gage and Axel (2014). While the authors reported the results of such an analysis, it encompassed the entire Twin Lakes database (197,845 recordings at the six sites).

Here, the soundscape power was derived from the 1440, 1-s recordings extracted from the original database and used in the present experiments. Each time of day was based on the average of 360 samples (12 samples × 10 days × 3 months). The soundscape power for 10 1-kHz frequency intervals (1 to 11 kHz) at the four times of day are shown in Fig. 6. Variations with time of day were negligible for frequency bands centered above 4 kHz, but were marked for the frequency bands centered at 1.5 and 2.5 kHz. The time of day could potentially be predicted from the differences in level between the 1-2 kHz and the 2-3 kHz bands, which were 0, 2.7, −1.6 and −0.8 dB for Dawn, Midday, Dusk and Night, respectively. Note that the two times of day with the largest values of the differences, Midday and Dusk, were also the two times giving the highest discrimination scores (left panel in Fig. 5), while the two times with the smallest differences, Dawn and Night, gave the lowest discrimination scores.

**Figure 6:**
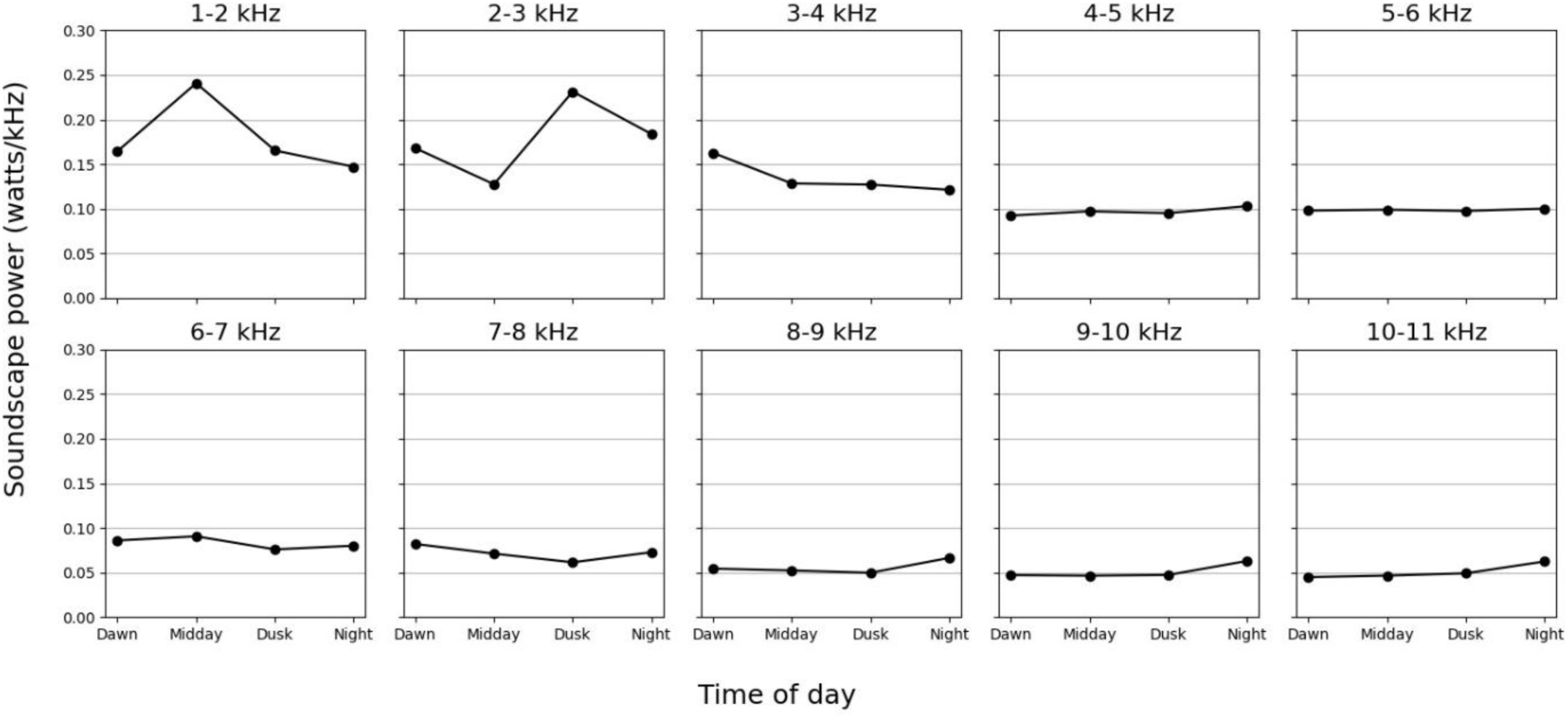
Spectral power as a function of the time of day for 10 1-kHz frequency intervals (1 to 11 kHz) computed over the 1440, 1-s recordings extracted from the original TL database. Each time of day corresponds to the average of 360 samples.

Performance was somewhat affected by the season, varying from 54% (Spring) to 60% (Fall) for UNF and from 49% (Spring) to 61% (Summer) for REC. A repeated-measures ANOVA of the discrimination scores with processing type (2 levels) and season (3 levels) as within-subjects factors showed no significant effect of processing type [*F*(1,8)=0.16; *p*=0.704] and a significant effect of season [*F*(2,16)=8.29; *p*<0.01]. The interaction between processing type and season was not significant [*F*(2,16)=2.38; *p*=0.12]. Spring was the season with the lowest score, for both conditions. This influence of seasonality on discrimination is consistent with a role of geophony and biophony cues, as these two factors are influenced by the season (Farina and Gage, 2017). However, Spring is typically associated with high biophony (the breeding season), but this season was associated with the lowest performance.

#### Id. Discrimination task with equalized rms level

In experiments Ia to Ic, natural level cues were preserved. Experiment Id assessed the contribution of level cues to the discrimination of changes in time of day by equating the rms level of all stimuli, as was done by Apoux *et al*. (2023). This was done only for condition UNF of Experiment Ib. Mean discrimination scores were 57.3% (SD = 6.1) for Experiment Ib and 56.6% (SD = 3.7) for Experiment Id; the difference was not significant (paired-samples *t*-test; *p*=0.64). These results indicate that level cues did not play a role in experiments Ia to Ic. It seems likely that they also did not influence the results of Apoux *et al*. (2023) for time of day discrimination.

## IV. EXPERIMENT II

The second experiment aimed to further explore the ability to discriminate time of day, using two additional databases. The primary objective was to evaluate whether the findings for TL would generalize to other biomes. A secondary goal was to investigate the potential influence of biophony. Apoux *et al*. (2023) suggested that discrimination may improve as acoustic activity and diversity increase. This hypothesis was indirectly tested by selecting biomes based on their expected biodiversity, informed by the “latitudinal gradient of diversity.” If biophony plays a significant role in the discrimination of times of day, performance should be similar for terrestrial biomes with comparable levels of biodiversity (biomes at similar latitudes), but differ for biomes with different levels of biodiversity (biomes at different latitudes).

Two biomes were selected from the database of the Hearbiodiv consortium (see Miller-Viacava *et al*., 2025). The first, HJ, was a temperate forest, sharing many features with TL, such as seasonal changes and layered vegetation structures. Both locations have mixed forests of deciduous and coniferous trees, though the ratio of these types differs: HJ is dominated by conifers. TL and HJ have similar latitudes (45.5401° and 46.6608°, respectively) but show notable differences in species composition. According to the latitudinal gradient of diversity, these biomes should have comparable levels of acoustic biodiversity. Thus, if biophony influences discrimination performance, no significant difference between TL and HJ would be expected in terms of overall performance.

The second biome, No, is a tropical forest, which shares fewer characteristics with TL than HJ does. Located closer to the Equator (4.0333°), No is expected to exhibit higher biodiversity than temperate forests, according to the latitudinal gradient of biodiversity. If biophony affects discrimination of time of day, performance should be higher for No than for either TL or HJ, due to the greater acoustic biodiversity of tropical forests.

Despite our efforts to select the most comparable biomes, notable differences remain between TL and HJ, as they belong to different continents. Thus, while the possibility that discrimination performance may differ between these two temperate forests cannot be excluded, it seems plausible that if biophony affects performance of the task, performance should be similar for TL and HJ and lower than for No.

### A. Methods

To avoid bias associated with previous experience of TL, 15 normal-hearing listeners (10 females) who did not participate in our previous studies took part in Experiment II. They were aged between 21 and 36 years (mean = 27 years; SD = 4.6 years).

Discrimination was tested in three sessions, each corresponding to one biome. The method was the same as for Experiment I. Stimuli were not altered in any way, except for the level adjustment described in the General Methods section. The three biomes were tested in random order across participants. Experiment II lasted about 2 hours (including breaks).

### B. Results

Figure 7 shows individual time of day discrimination scores for each biome. The scores ranged from 61 to 78% for HJ, from 61 to 84% for No, and from 49 to 69% for TL. Again, all scores were well above chance level (33%). The average scores were 69, 77, and 59% for HJ, No and TL, respectively. For most listeners, performance was best for No and worst for TL. Out of the 15 participants, only 3 showed a different pattern, where the highest score was for HJ.

**Figure 7:**
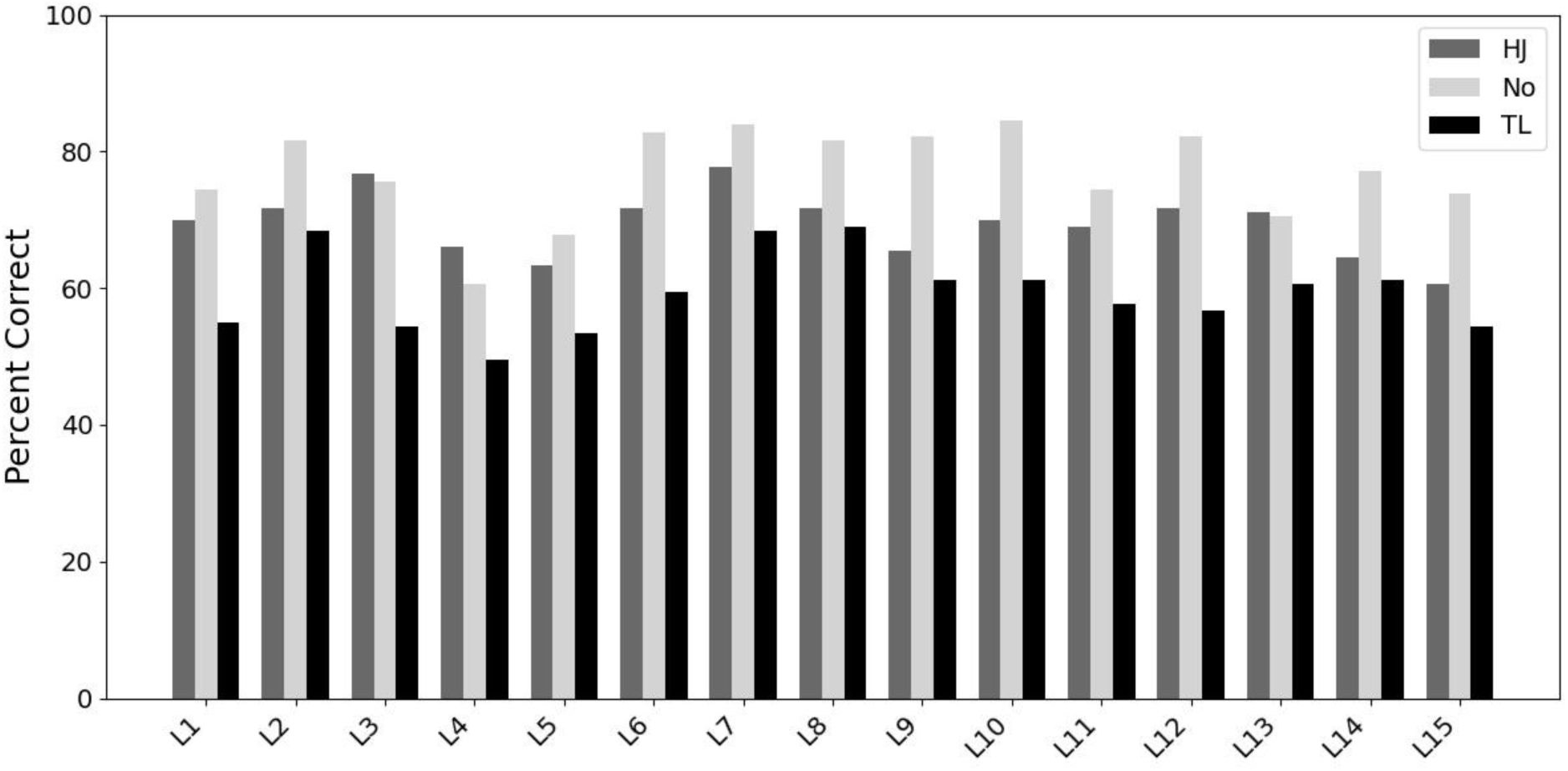
Individual data for Experiment II. Scores for discrimination of time of day are plotted for the three biomes: Haut Jura (HJ), Nouragues (No) and Twin Lakes (TL).

**Figure 8:**
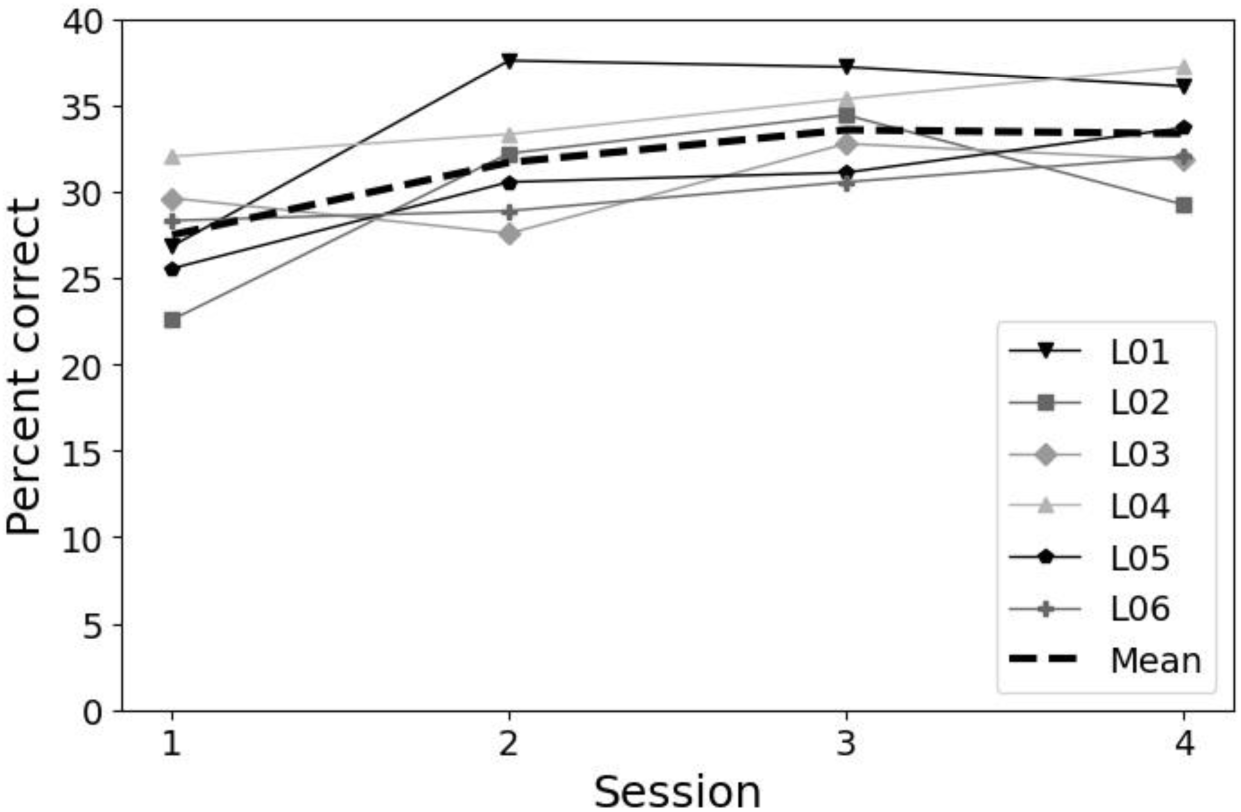
Individual data for Experiment III. Scores for discrimination of time of day are plotted as a function of training session. Session 1 was completed approximately one year before sessions 2 to 4, while the latter were completed within one hour. The dashed line shows the score averaged across all six participants. The chance level is 25% correct.

A repeated-measures ANOVA was conducted on the scores with Biome (3 levels) as within-subjects factor. The effect of Biome was significant [*F*(2,42)=33.89; *p*>0.001]. Post-hoc comparisons (Tukey HSD test) showed that all the scores were significantly different (all *p*<0.01) from each other, confirming the general pattern described above (No > HJ > TL). These findings are broadly consistent with an influence of the level of biophony activity, since scores were highest for the biome with the greatest species richness. However, the significant difference between HJ and TL is not consistent with the initial hypothesis based on the latitudinal gradient of diversity. This inconsistency may have occurred because biogeographical and ecological factors other than latitude can influence diversity levels, including altitude, vegetation and large-scale patterns of biodiversity distribution. For example, species richness tends to decrease with increasing elevation (Rahbek, 1995). The altitude difference of 1000 m on average between TL and HJ might have led to greater biodiversity for the latter. Additional work is necessary to estimate more accurately the level of biophony in each biome and to determine if that influences discrimination of time of day.

## V. EXPERIMENT III

Experiment I showed only very limited abilities to identify time of day. This may have occurred because the participants did not have sufficient experience in associating the acoustic properties of the scenes with time of day. Experiment III explored the effect of training on the ability to identify time of day. The method was similar to that for Experiment Ia (identification task), except that in Experiment III listeners were provided with the correct response (i.e. the time of day) after each trial, rather than being told that the response was correct or incorrect.

### A. Method

Six normal-hearing listeners (3 females) took part. They had all participated in Experiment I. Therefore, they were familiar with the stimuli, and they had already completed the identification task once with limited feedback. The time interval between the two experiments was approximately one year. Listeners were aged between 21 and 37 years (mean = 30.2 years; SD = 5.5 years).

### B. Results

Figure 8 presents the individual identification scores and the average score (dashed line) for the initial session with limited feedback, and for the three subsequent sessions with precise feedback, in chronological order. The dashed line represents the identification scores averaged across all six participants. The mean scores were 28% for session 1, and 32, 34, and 33% for sessions 2, 3, and 4, respectively. A repeated-measures ANOVA on the scores with Session (4 levels) as within-subjects factor showed a significant main effect [*F*(3,20)=4.91; *p*<0.05]. Post-hoc comparisons (Tukey HSD test) showed that the score for the initial session was only significantly different from scores for sessions 3 and 4 (all *p*<0.05). Thus, there was a slight improvement in performance with more precise feedback, with an increase of 4-percentage points between session 1 and session 2, followed by a 2-percentage point increase from session 2 to session 3. However, no further improvement from session 3 to session 4. The marginal improvement in scores could be attributed to the precise feedback provided during sessions 2 through 4, though it is also possible that the participants’ prior experience in the discrimination experiments (Exp. I), which involved over 8 hours of listening per participant, played a role.

Even with precise feedback and extensive practice, mean scores were only a little above the chance level of 25% correct. It seems that the acoustic information in natural soundscapes is not sufficient to allow precise identification of time of day, at least from 1-s samples.

## VI. DISCUSSSION

### A. Discrimination of time of day for natural soundscapes

The three experiments confirmed that human listeners can discriminate recordings of natural soundscapes based on the time of day with reasonably high sensitivity (d′ scores ranged between 1.6 and 3.4 for the unprocessed conditions), consistent with previous work (Apoux *et al*., 2023). Importantly, the results show that good discrimination is not restricted to recordings from a specific habitat, since high sensitivity was observed for three habitats differing in terms of location and/or latitude. Discrimination performance seems to be reasonably consistent both within and across habitats. Apoux *et al*. (2023) reported discrimination scores ranging across participants from 55 to 68% correct, averaged across four habitats in the Sequoia National Park (SEKI), an area located in the southern Sierra Nevada, east of Visalia, California (USA) that may be classified as a Mediterranean biome. In the present study, performance ranged across participants from 60.6 to 77.8%, 60.6 to 84.4% and 49.4 to 68.9% for HJ, No and TL, respectively. The present results are also consistent with previous conclusions that little to no exposure to a specific soundscape is needed to discriminate time of day in that soundscape.

Also consistent with Apoux *et al*. (2023), the results of Experiments Ib and Ic show that stimulus variability associated with changes in season does not strongly affect the ability to discriminate time of day. This is surprising, since animals and insects are known to be more, if not only, active and vocal during reproduction seasons, typically spring at high and low latitudes. Assuming that biophony plays a role in natural soundscape discrimination, higher performance might have been expected for locations and seasons where biophony is high. The present data are not consistent with this hypothesis, as spring was the season for which discrimination performance was lowest overall.

Although performance differed across time of day, this effect was small. According to the analysis of Gage and Axel (2014) and the biophony hypothesis, it was assumed that time of day associated with the dawn or dusk choruses would be easier to discriminate, as listeners could rely more effectively on vocalisations to perform the task. The present data, however, show that this was not the case. Dusk was better discriminated than the other times of day for the unprocessed condition, but dawn was not. One may also have expected night to be relatively well discriminated, as it is associated with animals that are not heard during the day. In the present experiments, however, night was the time of day associated with the poorest performance both when it was a target and when it was a standard. While further work is warranted to determine the cues used by listeners to discriminate time of day, it is apparent that there is no simple relationship between the quantity or the quality of the biophonic content in a sample and its discriminability.

It should be remembered that discrimination of time of day was measured using a 3-alternative forced-choice (3AFC) procedure in which each trial contained one target (e.g. a sample drawn from dawn) and two standards (e.g. both samples drawn from dusk). However, unlike in a conventional 3AFC procedure, the two standards within a trial were not identical; both were drawn from the same time of day, but they were different samples from that time of day. Thus, the task could not be performed by detecting any difference at all between the target and the standards, since the two standards within a trial would have differed. Rather, to perform the task reasonably well, some feature of the target had to be detected that usually distinguished it from the two standards. This may account for the finding that while discrimination performance was reasonably good, it was never perfect, or even close to perfect.

### B. Identification vs. discrimination of time of day in natural soundscape

One goal of the present work was to investigate both discrimination and identification of the time of day of natural soundscapes. Identification was found to be poor (about 28% correct), but above chance, before extensive training and using only imprecise feedback (“correct” or “wrong”). Identification improved following training with more precise feedback (being told the correct time of day), but remained relatively poor, at about 33%, only a little above chance performance of 25%. This low performance contrasts with the reasonably good performance for the discrimination tasks. The discrimination results indicate that natural soundscapes differ according to the time of day and that human listeners are sensitive to these differences. However, the identification results show that listeners are not able to consistently associate those differences with the correct label: “dawn”, “dusk”, etc.).

This difficulty may in part be attributed to the complexity and the high variability of the stimuli. Not only do natural soundscapes potentially contain a multitude of sound sources, but some of these sources are not present in every 1-s sample. Longer sample durations might have alleviated this problem by providing a more consistent set of cues in each sample. Another factor to bear in mind is that on a given trial of the discrimination task, the target time of day had to be discriminated only from one other time of day (the time of day of the two standards), while in the identification task the time of day had to be identified from four possibilities.

### C. Cues involved in discrimination of time of day for natural soundscape

The present work also addressed the cues used by the participants in the discrimination experiments, and particularly the role of biotic information. The audio-filtering experiment (Experiment Ib) showed that the best performance was obtained using the unfiltered stimuli. Performance worsened as the cutoff frequency decreased in the LP condition and as the cutoff frequency increased in the HP condition. However, reasonably good performance was still achieved when the stimuli were lowpass filtered at 500 Hz or highpass filtered at 4000 Hz. Thus, usable information was available over a wide audio-frequency range. The average crossover frequency was around 2.2 kHz, slightly higher than what was observed for habitat discrimination (i.e., 1.8 kHz), but the similarity between the two values suggests that the same conclusion may apply to the present data. As a minimum, it may be assumed that the greater weight attributed by the participants to acoustic cues in the mid-frequency range may reflect a role played by biophony (Gage and Axel, 2014).

The envelope-filtering experiment (Experiment Ic) failed to show a contribution of specific temporal envelope cues to time of day discrimination using vocoded stimuli. Removing either the low- or high-rate temporal-envelope information affected time of day discrimination only slightly and non-significantly. It appears that cues sufficient to give reasonable discrimination were available at both low and high envelope modulation rates. Performance was only a little worse for the noise-vocoded stimuli than for the REC stimuli, suggesting that near-intact TFS information and fine spectral detail were not critical for discrimination performance. Because some factors, such as sample duration, may have limited the use temporal cues, it is not possible from the present data to draw a clear conclusion and more work is warranted to exclude the role of amplitude modulations in this task, especially in a more ecological context.

It was possible that participants would use differences in level across samples to perform the task. For instance, rain or wind may have been consistently present at one time of day, which would have increased the level of the corresponding samples. However, the results of Experiment Id were not consistent with this possibility, since equalizing the rms level had no effect on time of day discrimination. It should be noted, however, that equating the rms level across samples would not have equated loudness across samples, although it probably reduced loudness differences across samples. Further work is needed to establish whether loudness differences can be used to discriminate time of day in natural soundscapes.

Animal vocalizations have been associated with slow temporal modulations in natural soundscapes (Gerhardt and Huber, 2002; Singh and Theunissen, 2003; Marler and Slabbekoorn, 2004; Catchpole and Slater, 2008; Di Tullio *et al*., 2024) and one hypothesis of the present work was that biophony would play a role in time of day discrimination. The present data were not consistent with this hypothesis, since highpass filtering of the envelopes of vocoded stimuli to remove slow modulations had little effect on discrimination performance. It should be noted, however, that the database used in Experiment Ic came from a temperate forest and such habitats are not especially rich in animal vocalizations. Another factor that may have restricted the use of biophony in the present experiments is the 1-s duration of the stimuli. Combined with the relatively limited amount of animal vocalization, this short duration resulted in many samples containing no biophony. Informal listening to the entire database confirmed that this was the case. As a result, our participants may not have been able to reliably base their decisions on identifiable biotic events, such as bird vocalizations or insect chirps. They may instead have relied on global acoustic attributes of the soundscape.

The results of Experiment II provided some evidence supporting a role of biophony in time of day discrimination. They showed better performance for the biome with higher species richness (the tropical forest) than for the less rich biomes. As mentioned previously, however, this is very indirect evidence, and other factors may have contributed to this advantage. Clearly, additional work is needed to determine the role of biophony in time of day discrimination. Overall, the present data suggest that fine spectral details, temporal envelope cues at specific frequencies, and TFS cues are not critical for discriminating time of day. Consistent with previous work, it is suggested that human listeners rely primarily on long-term spectral cues.

### D. Use of global attributes of soundscapes and textures

Previous work has shown that the diel cycle is strongly associated with variations in animal vocalizations and activity (e.g., Gage and Axel, 2014). Accordingly, it was assumed that biotic cues would provide information for roughly determining the time of day and that participants would use this information. As mentioned above, however, the data did not support a role of biophony in time of day discrimination.

In their study, McMullin *et al*. (2024) aimed to compare the contribution of detail-oriented and global processes in identifying seasonality, among other aspects. Detail-oriented processes are related to the segregation and identification of individual sources in the scene. The categorization of a scene based on such processes would rely on the identification of single sound sources that are assumed to be stereotypical of such an acoustic environment. For instance, one would categorize a soundscape including cow sounds as a farm scene. This has been demonstrated in vision by showing, for instance, that participants are able to identify scenes even from a short presentation of a single object (Wiesmann and Võ, 2023). Global processes are related to the extraction of information about the constancy, structure, or function of a scene (Greene and Oliva, 2009a, 2009b; Ross and Oliva, 2010). They do not require identification of individual sound sources. This “high-level” information includes global properties of the scene such as navigability (the scene is clear from obstacles, e.g., a meadow, vs. it contains many obstacles, e.g., a forest) or naturalness (the scene is natural, e.g., a forest, vs. a man-made environment, e.g., a parking lot). McMullin *et al*. (2024) suggested that human listeners extract global properties or attributes from auditory scenes. Global properties are believed to be independent of the identification and segregation of specific sources or objects within the scene. For example, listeners may not need to identify the wind in the trees to establish that a recording was made in a closed environment such as a forest. Similarly, listeners may not need to identify specific animal vocalizations to extract some information about the time of day from the global sound of vocalizations. These global attributes alone may provide important information about the scenes and therefore support auditory discrimination and identification. The results of this study suggest that biophony information is not necessary for extracting global features such as seasonality. If similar global processes are involved in identifying and discriminating time of the day, this may partially account for the limited impact of biophony.

Overall, our results suggest that human listeners monitor the gross acoustic features (potentially, long-term spectral cues) of sound to discriminate diel changes. This is consistent with the proposal of Fay (2009) that scattering, reverberation and non-identifiable contributions of biophony and geophony inherently contained in the “ambient noise field” provide cues that could be used by human and non-human observers to obtain information about environmental content and its daily or seasonal variations. In other words, “the ambient noise field itself has the potential to be exploited for imaging the environment” (Fay, 2009). More work is warranted to assess the exact information conveyed by the ambient noise field of natural settings and its use in daily tasks by human listeners.

## VII. CONCLUSIONS

1) Human listeners can discriminate short samples of natural soundscapes that differ only in broadly defined time of day. This was found to be true for three soundscapes recorded from different biomes. The soundscape recorded closer to the equator led to the best discrimination performance, perhaps partly because of greater biodiversity.
2) Human listeners struggled to identify time of day based on 1-s samples of the soundscape. While performance was significantly above chance level, it remained very poor even after one hour of training with feedback. This finding suggests that while human listeners are sensitive to acoustic variations related to differences in time of day, they cannot consistently label these variations.
3) Despite inconclusive data, it is suggested that biophony – the collective sound produced by animal vocalizations – did not play a critical role in time of day discrimination, probably because this cue was often inconsistent across samples. Given the short duration of the samples used here, it is possible that longer durations will lead to better performance via a greater contribution of biotic information.
4) Level, fine spectral, temporal envelope and TFS cues were all tested as potential candidates for supporting time of day discrimination. Surprisingly, none of these cues was essential for performance of the task. Consistent with previous work, it is suggested that human listeners based their decisions on gross spectral cues.

## ACKNOWLEDGMENTS

This work was supported by ANR-17-EURE-0017, ANR-20-CE28-0011 and ANR AUDIECO. The work conducted in the Risoux forest was supported by the Parc Naturel Régional du Haut-Jura, which received funding from the Région Bourgogne-Franche-Comté, the Région Auvergne-Rhône-Alpes, and the DREAL Bourgogne-Franche-Comté. The work conducted in the Nouragues forest was supported by Labex CEBA, which received funding from Investissements d’Avenir ANR-10-LABX-25-01. Sylvain Haupert and Frédéric Sèbe are warmly thank for their help in collecting the data. The authors would like to thank Syndicat National des Audioprothésistes for its continued support.

## Footnotes

1 These times were computed using the Matlab implementation by Droste (2022) of the sunrise/sunset and solar position calculations developed by the USA National Oceanic and Atmospheric Administration, based on equations from Astronomical Algorithms by Meeus (1991). With this implementation, the sunrise and sunset times were corrected for atmospheric refraction effects so that they were closer to the real time when sunlight changed; the daylight savings time change was also taken into account when needed as UTC corrections for the corresponding months (UTC is an input for the Matlab function).

## REFERENCES

Alcocer, I., Lima, H., Sugai, L. S.M., and Llusia, D. (2022). “Acoustic indices as proxies for biodiversity: a meta-analysis,” Biol. Rev. Camb. Philos. Soc., 97, 2209–2236. 10.1111/brv.12890

Apoux, F., Miller-Viacava, N., Ferrière, R., Dai, H., Krause, B., Sueur, J., and Lorenzi, C. (2023). “Auditory discrimination of natural soundscapes,” J. Acoust. Soc. Am., 153, 2706–2723. 10.1121/10.0017972

Catchpole, C.K., and Slater, P.J.B. (2008). Bird song: Biological themes and variations (Cambridge University Press, Cambridge), pp. 1–348.

de Baudouin, A., Couprie, P., Michaud, F., Haupert, S., and Sueur, J. (2023). “Similarity visualization of soundscapes in ecology and music,” Front. Ecol. Evol., 12, 1–13. 10.3389/fevo.2024.1334776

Di Tullio, R.W., Wei, L., and Balasubramanian, V. (2024). “Slow and steady: auditory features for discriminating animal vocalizations”, bioRxiv 2024.06.20.599962. 10.1101/2024.06.20.599962

Farina, A. and Gage, S.H. (2017). Ecoacoustics: The ecological role of sounds (John Wiley & Sons, Hoboken), pp. 1–336.

Fay, R. (2009). “Soundscapes and the sense of hearing of fishes,” Integr. Zool., 4, 26–32. 10.1111/j.1749-4877.2008.00132.x

Frijters, J.E.R. (1979). “Variations of the triangular method and the relationship of its unidimensional probabilistic models to three-alternative forced-choice signal detection theory models,” Brit. J. Math. Stat. Psychol., 32, 229–241. 10.1111/j.2044-8317.1979.tb00595.x

Frijters, J.E.R., Kooistra, A., and Vereijken, P.F.G. (1980). “Tables of d’ for the triangular method and the 3-AFC signal detection procedure,” Percept. Psychophys., 27, 176–178. 10.3758/BF03204306

Gage, S.H., and Axel, A.C. (2014). “Visualization of temporal change in soundscape power of a Michigan lake habitat over a 4-year period,” Ecol. Inform., 21, 100–109. 10.1016/j.ecoinf.2013.11.004

Gaston K.J. (2000). “Global patterns in biodiversity,” Nature, 405, 220–227. 10.1038/35012228

Gerhardt, H.C., and Huber, F. (2002). Acoustic communication in insects and anurans: Common problems and diverse solutions (The University of Chicago Press, Chicago, IL), pp. 1–542.

Gil, D., Llusia, D. (2020). The Bird Dawn Chorus Revisited. In: Aubin, T., Mathevon, N. (eds) Coding Strategies in Vertebrate Acoustic Communication. Animal Signals and Communication, vol 7. Springer, Cham. 10.1007/978-3-030-39200-0_3

Glasberg, B.R., and Moore, B.C.J. (1990). “Derivation of auditory filter shapes from notched-noise data,” Hear. Res. 47, 103–138.

Greene, M.R., and Oliva, A. (2009a). “The briefest of glances: The time course of natural scene understanding,” Psychol. Sci., 20, 464–472. 10.1111/j.1467-9280.2009.02316.x

Greene, M.R., & Oliva, A. (2009b). “Recognition of natural scenes from global properties: Seeing the forest without representing the trees,” Cogn. Psychol., 58, 137–176. 10.1016/j.cogpsych.2008.06.001

Grinfeder, E., Lorenzi, C., Haupert, S., and Sueur, J. (2022). “What do we mean by “soundscape”? A functional description,” Front. Ecol. Evol., 10, 894232 10.3389/fevo.2022.894232

Gygi, B., Kidd, G., and Watson, C. (2004). “Spectral-temporal factors in the identification of environmental sounds,” J. Acoust. Soc. Am., 115, 1252–1265. 10.1121/1.1635840

Hillebrand, H. (2004). “On the generality of the latitudinal diversity gradient,” Am. Nat., 163, 192–211. 10.1086/381004

Keidser, G., Naylor, G., Brungart, D.S., Caduff, A., Campos, J., Carlile, S., Carpenter, M.G., Grimm, G., Hohmann, V., Holube, I., Launer, S., Lunner, T., Mehra, R., Rapport, F., Slaney, M., and Smeds, K. (2020). “The Quest for Ecological Validity in Hearing Science: What It Is, Why It Matters, and How to Advance It,” Ear. Hear., 41, 5–19. 10.1097/AUD.0000000000000944

Krause, B. (2016). Wild soundscapes: discovering the voice of the natural world (Yale University Press, New Haven), pp. 1–240.

Krause, B., Gage, S.H., and Joo, W. (2011). “Measuring and interpreting the temporal variability in the soundscape at four places in Sequoia National Park,” Landsc. Ecol., 26, 1247–1256. 10.1007/s10980-011-9639-6

Loo, Y.Y., Lee, M.Y., Shaheed, S., Maul, T., and Clink, D.J. (2025). “Temporal patterns in Malaysian rainforest soundscapes demonstrated using acoustic indices and deep embeddings trained on time-of-day estimation,” J. Acoust. Soc. Am., 157, 1–16. 10.1121/10.0034638

Lorenzi, C., Apoux, F., Grinfeder, E., Krause, B., Miller-Viacava, N., and Sueur, J. (2023). “Human auditory ecology: Extending hearing research to the perception of natural soundscapes by humans in rapidly changing environments,” Trends in hearing, 27, 23312165231212032. 10.1177/23312165231212032

Marler, P., and Slabbekoorn, H. (2004). Nature’s music: The science of birdsong (Elsevier Academic Press, San Diego, CA), pp. 1–513.

McMullin, M.A., Kumar, R., Higgins, N.C., Gygi, B., Elhilali, M., and Snyder, J.S. (2024). “Preliminary Evidence for Global Properties in Human Listeners During Natural Auditory Scene Perception,” Open mind: discoveries in cognitive science, 8, 333–365. 10.1162/opmi_a_00131

Miller-Viacava, N., Apoux, F., Ferriere, R., Friedman, N.R., Mullet, T.C., Sueur, J., Willie, J., and Lorenzi, C. (2025). “Modulation statistics of natural soundscapes,” bioRxiv 2025.03.04.638661. 10.1101/2025.03.04.638661

Moore, B. C. J. (2014). Auditory Processing of Temporal Fine Structure: Effects of Age and Hearing Loss (World Scientific, Singapore), pp. 1–182.

Moore, B. C. J., Füllgrabe, C., and Sęk, A. (2009). “Estimation of the center frequency of the highest modulation filter,” J. Acoust. Soc. Am. 125, 1075–1081. 10.1121/1.3056562

Neuhoff, J. (2004). Ecological Psychoacoustics (Elsevier Academic Press, Cambridge, MA), pp. 1–350.

Pijanowski, B.C., (2024). Principles of soundscape ecology: Discovering our sonic world (The University of Chicago Press, Chicago, IL), pp. 1–456.

Pijanowski, B.C., Villanueva-Rivera, L.J., Dumyahn, S.L., Farina, A., Krause, B.L., Napoletano, B.M., Gage, S.H., and Pieretti, N. (2011). “Soundscape ecology: the science of sound in the landscape,” Bioscience, 61, 203–216. 10.1525/bio.2011.61.3.6

Rahbek, C. (1995). “The elevational gradient of species richness: a uniform pattern?,” Ecography, 18, 200–205. 10.1111/j.1600-0587.1995.tb00341.x

Rolland, J., and Freeman, B.G. (2023). “The latitudinal diversity gradient,” Evol. Biol. 10.1093/obo/9780199941728-0144

Ross, M.G., and Oliva, A. (2010). “Estimating perception of scene layout properties from global image features,” J. Vision, 10, 1–25. 10.1167/10.1.2

Schmuckler, M.A. (2001). “What is ecological validity? A dimensional analysis,” Infancy, 2, 419–436. 10.1207/s15327078in0204_02

Shafiro, V. (2008). “Identification of environmental sounds with varying spectral resolution,” Ear. Hear., 29, 401–420. 10.1097/AUD.0b013e31816a0cf1

Singh, N.C., and Theunissen, F.E. (2003). “Modulation spectra of natural sounds and ethological theories of auditory processing,” J. Acoust. Soc. Am., 114, 3394–3411. 10.1121/1.1624067

Sueur, J., and Farina, A. (2015). “Ecoacoustics: the ecological investigation and interpretation of environmental sound,” Biosemiotics, 8, 493–502. 10.1007/s12304-015-9248-x

Thoret, E., Varnet, L., Boubenec, Y., Ferriere, R., Le Tourneau, F.-M., Krause, B. and Lorenzi, C. (2020). “Characterizing amplitude and frequency modulation cues in natural soundscapes: A pilot study in four habitats of a biosphere reserve,” J. Acoust. Soc. Am., 147, 3260–3274. 10.1121/10.0001174

Varnet, L., Ortiz-Barajas, M. C., Erra, R. G., Gervain, J., and Lorenzi, C. (2017). “A cross-linguistic study of speech modulation spectra,” J. Acoust. Soc. Am., 142, 1976–1989. 10.1121/1.5006179

Versfeld, N.J., Dai, H., and Green, D.M. (1996). “The optimum decision rules for the oddity task,” Percept. Psychophys., 58, 10–21. 10.3758/bf03205470

Welch, P. (1967) “The use of Fast Fourier Transform for the estimation of power spectra: A method based on time averaging over short, modified periodograms,” IEEE Transactions on Audio and Electroacoustics, 15, 70–73. 10.1109/TAU.1967.1161901

Wiesmann, S.L., and Võ, M.L. (2023). “Disentangling diagnostic object properties for human scene categorization,” Sci. rep., 13, 5912. 10.1038/s41598-023-32385-y

Willig, M., Kaufman, D., and Stevens, R. (2003). “Latitudinal gradients of biodiversity: pattern, process, scale, and synthesis,” Annu. Rev. Ecol. Evol. Syst., 34, 273–309. 10.1146/annurev.ecolsys.34.012103.144032

